# Mimicry in motion: A grasshopper species that looks like, and moves like, a sympatric butterfly

**DOI:** 10.1101/2023.06.04.543626

**Authors:** Karl Loeffler-Henry, Changku Kang, Jeff W. Dawson, Thomas N. Sherratt

**Affiliations:** Department of Biology, Carleton University; Ottawa, Ontario, Canada; Department of Agricultural Biotechnology, Seoul National University; Seoul, South Korea; Research Institute of Agricultural and Life Sciences, Seoul National University; Seoul, South Korea

## Abstract

Flash displays describe the sudden revealing of a conspicuous signal by an otherwise cryptic animal as it is fleeing from predators. One hypothesis for flash displays is that they have evolved to mislead predators into adopting an inappropriate search strategy following pursuit. Thus, if predators believe that the fleeing prey item will also be conspicuous when settled, then they will tend to give up searching sooner if they fail to find it. Naturally, if this hypothesis is correct then one might expect that the value of the flash display would cease once predators learn the flasher’s true resting appearance. However, even if predators are aware that the resting appearance of the signaller could be cryptic, flash displays will still be selected for if the flasher resembles another species that maintains its conspicuous appearance throughout. Motivated by this insight, we quantified the degree of chromatic and behavioral resemblance between the Carolina grasshopper (*Dissosteira carolina*) and mourning cloak butterfly (*Nymphalis antiopa*), a sympatric species with which it is commonly confused. The close similarity in both morphology and flight behavior (in the field and lab) between these distantly related species strongly suggests that they are in a mimetic relationship. It is unclear why this mimicry has evolved. One explanation is that mimicry has arisen through selection to maintain the efficacy of the flash display through search disruption even after predator learning. Another possibility is that the grasshopper has evolved to resemble the butterfly because the latter is harder to catch (evasive mimicry). However, we argue that pursuit and search disruption are entirely complementary explanations for the mimetic resemblance because they interfere with predation at two different stages of the attack sequence.

## Introduction

Some animals are cryptic at rest but expose a burst of colour as they flee from would-be predators, returning to their cryptic appearance as they settle [1, 2]. Examples of this “*flash behaviour*” are taxonomically widespread, having been described in all five classes of vertebrate as well as insects and cephalopods [3]. Morphologically, flash displays are typically generated by the exposure of high contrast conspicuous colours - such as the hindwings of insects, and the flight feathers of birds - that are only revealed during movement [1]. Alternatively, a conspicuous signal may be revealed through the use of chromatophores, as seen in some fleeing cephalopods [1, 4].

Several hypotheses have been proposed to explain the adaptive significance of flash behaviour. In some cases, the signal may serve to convey information to conspecifics [5]. Alternatively, the sudden exposure of a previously hidden conspicuous signal may startle a predator on deployment, buying time for the signaller to escape [3, 6]. Dynamically changing colors while in motion may also lower the probability that a prey is captured [7-9]. Another proposed function of flash behaviours is that it misleads would-be predators into believing the “flasher” is always conspicuous in appearance, which hinders the predator’s subsequent search for it [1, 3, 10]. This phenomenon has recently been demonstrated to be quantitatively plausible using Bayesian search theory [11]. Specifically, the more conspicuous a prey item is believed to be, the sooner a predator should give up searching for it in an area where it appears to have settled; so cryptic prey giving the impression of being conspicuous will have a higher rate of survival than those that do not. Recent experimental studies have also supported this ‘misdirection’ hypothesis, albeit with artificial prey. In particular, a computer-based experiment found that human participants were more likely to give up looking for prey that were a conspicuous color when in motion and cryptic when sedentary, compared to those prey that were cryptic throughout [12]. Further mechanistic support has come from a follow-up computer experiment which showed that flash displays in computer-generated prey only yielded a significant anti-predation benefit if the prey initiated their escape before the predator could observe the prey’s resting appearance [13].

The above experiment highlights a weakness of the anti-predator strategy: if the flashing species is commonly encountered, it is likely that predators will eventually learn to recognize that the displays are generated by a species that is cryptic at rest, undermining its benefits. On reflection however, if the flash display were to resemble another organism that retains its conspicuous appearance at rest, then one might expect it would reduce the rate at which predators make the association. Indeed, in the longer term even if the predator was completely aware of the mimicry, it may still give up looking sooner compared to a case where it knew the signaller was cryptic at rest, due to the increased uncertainty. A simple mathematical model which extends the model of Sherratt & Loeffler-Henry [11] to allow for mimicry supports this reasoning (see Appendix A, Supporting Information).

One insect species that is cryptic at rest but displays conspicuous colour patterns in flight is the Carolina grasshopper *Dissosteira carolina* [11]. Both males and females of this species exhibit the same black hindwings with yellow margins during flight, suggesting that the trait is not the product of sexual selection. Cooper [14] noted that with its sudden change in colour and movement, this grasshopper species becomes difficult to relocate when settled. The illusion may be reinforced by the fact that both the rapid takeoffs and landings of *D. carolina* will be perceived by a range of observers, including birds, to arise almost instantaneously [15]. Surprisingly however, one aspect of the flight behaviour of this species that was not mentioned in either of the above studies is the remarkable similarity of *D. carolina* to the sympatric mourning cloak butterfly *Nymphalis antiopa* (see Fig 1). Web-based descriptions for the species (for example [16-20]) repeatedly suggest that the colourful wings, large size and lazy flight of the grasshopper frequently lead it to be mistaken for the mourning cloak. Indeed, the seasoned entomologist John Acorn [21] quipped that he had “*been mistaking them for mourning cloaks since I was a kid*”. Naturally, while accounts by experienced field naturalists are indicative, first impressions can be misleading: direct quantitative evaluations of the extent of morphological and behavioral similarity are required to test the validity of these assertions, especially when potential predators such as birds can discriminate colour in the UV range of the spectrum [22].

**Fig 1.**
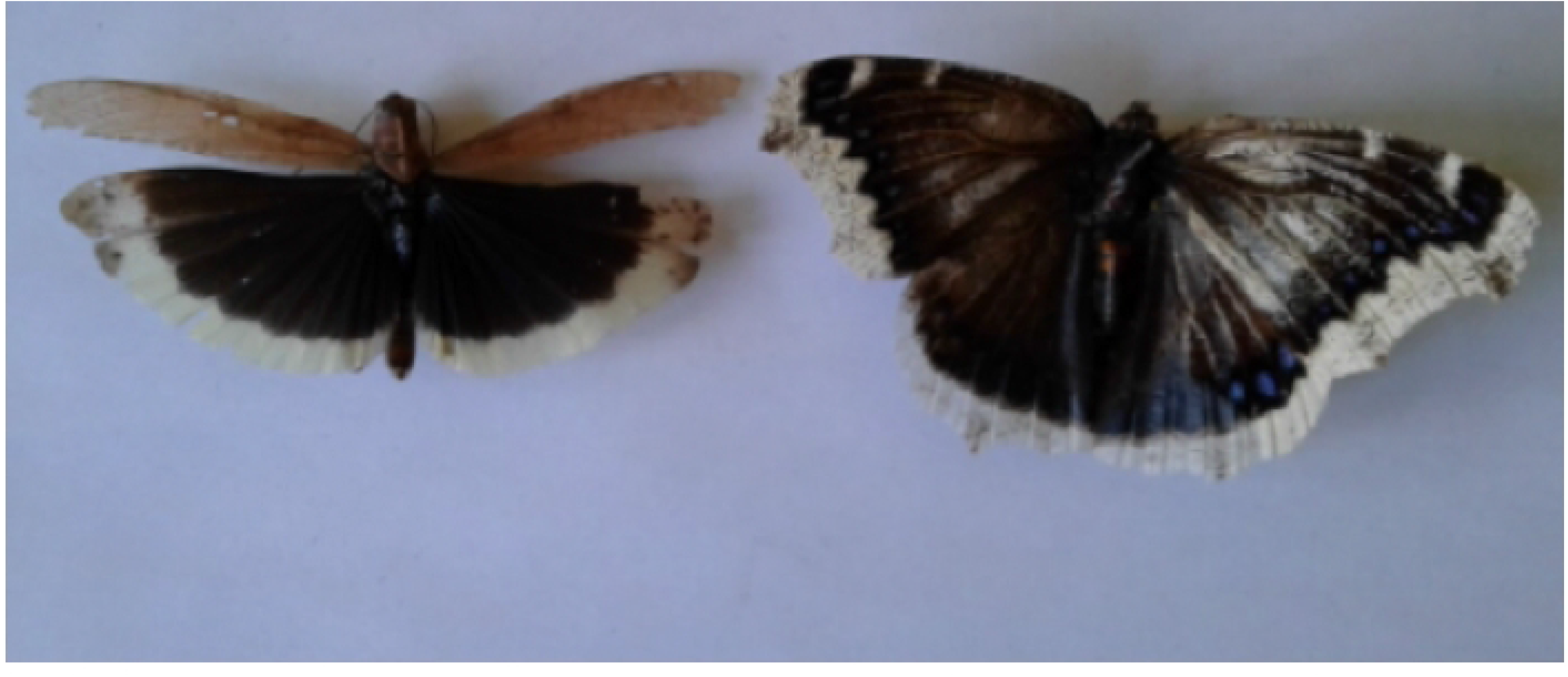
The wings (dorsal views) of the Carolina grasshopper *Dissosteira carolina* (left) and that of its potential sympatric model, the mourning cloak *Nymphalis antiopa* (right). Note the similarity (to human eyes) in their colour pattern. Photo credit: KL-H.

To test the possibility that the Carolina grasshopper and mourning cloak butterfly are in a mimetic relationship we determined their absolute and relative similarity (compared to other Orthoptera and other Lepidoptera) not just in terms of their visual appearance to would-be predators, but also in their flight behaviour. The visual appearance of the grasshopper and butterfly were compared assuming a bird visual model which allows for the possibility of colour discrimination in the UV range of the spectrum. The flight behaviour of each of our species was quantified in two complementary ways. First, detailed recordings of the flight of the Carolina grasshopper and mourning cloak butterfly and several other co-occurring orthopteran and lepidopteran species (selected on the basis of their relative abundance and sympatry) were made in a wind tunnel designed for insect flight. This approach allowed fine-scale measurements of flight behaviour of the insect species under a controlled setting. However, tethering individuals is clearly artificial. Therefore, to complement this work we recorded the behaviour of the Carolina grasshopper, mourning cloak and several other co-occurring insect species in the field. From these videos, we were able to extract quantitative metrics of flight behaviour under natural conditions. Evidence that *D. carolina* and *N. antiopa* not only share a morphological resemblance but also move in ways that are more similar than any other species pair would lend quantitative support to the hypothesis that these species are in a mimetic relationship.

## Methods

### Colour comparison of D. carolina and N. antiopa

For our colour comparison, we photographed two specimens of *D. carolina* (collected in Ottawa, ON, Canada) and two specimens of *N. antiopa* (held at the Natural History Museum London, originally collected in Rennes, France). The photographs we made using a full-spectrum converted Nikon D7000 camera equipped with Jenoptik UV-VIS-IR 60 mm 1:4 APO Macro lens. We used two lens filters to capture the human-visible (400 – 680 nm wavelength; Baader UV/IR cut filter, Mammendorf, Germany) and ultraviolet (320 – 380 nm; Baader U-Venus filter) range respectively. For *N. antiopa*, we used two bulbs that were placed next to each other for lighting the specimens: a True-light LED 12W bulb (CRI index 98; TRUE-LIGHT, Frankfurt, Germany) and a 3000K tungsten-halogen lamp (150W, Long Life Lamp Company, Harrow, UK). For *D. carolina*, we used a 175W metal-halide lamp (Dongsung, Namyangju, South Korea). Both lighting conditions emitted both the ultraviolet and human-visible range of light. We placed a 99% reflectance standard (WS- 1- SL, Labsphere, NH, USA) next to the specimens for later colour calibration. We used a constant camera setting (F8, ISO400) except for the shutter speed which was varied to avoid saturation (i.e. the pixel values of the white standard exceed the maximum pixel value) [23] that would otherwise arise in slightly under-exposed photos. All images were taken in raw format.

The raw images were analysed using the micaToolbox implemented in ImageJ 1.52a (open source program, National Institute of Health, USA) [24]. To examine whether the wing colours of *D. carolina* and *N. antiopa* look similar to the vision of their likely natural visual predators (birds), the raw images were converted to cone-catch images using a receptor-noise model that assumed blue tit (*Cyanistes caeruleus*) visual system [25]. Then we estimated the chromatic (derived from single cone catches) and achromatic (derived from double cone catches) just noticeable differences (JNDs). We selected and measured three random regions of each wing region using the free-hand selection tool and compared the cone catches between the two species. We used the average cone catch value of two specimens for each colour region to yield a single JND value. We assumed the Weber noise for long wavelength photoreceptors as 0.05, and the relative proportion of the different cone types in the retina as 1: 1.9: 2.7: 2.7 for the ultraviolet-sensitive, violet-sensitive, short-wavelength-sensitive, and long-wavelength-sensitive cone cells [25, 26].

### Wing beat characteristics of species in the laboratory

Laboratory observations of the flight characteristics of seven insect species were made from September 2017 to September 2019. Most insect specimens were obtained from the field at various sites around Ottawa, Ontario, Canada. These specimens were the grasshoppers: Carolina grasshopper *Dissosteira carolina* (*n* = 7)*, Melanoplus femurrubrum* (*n* = 7) and *M. bivittatus* (*n* = 6) along with the mourning cloak butterfly *Nymphalis antiopa* (*n* = 7) and the cabbage white butterfly *Pieris rapae* (*n* = 5). Specimens of two additional species (the grasshoppers *Locusta migratoria* (*n* = 7) and *M. sanguinipes* (*n* = 7) were obtained from Carleton University laboratory cultures. The *L*. *migratoria* culture has been maintained at Carleton University for 18 years, while the *M. sanguinipes* culture has been maintained at Carleton for 6 years (in turn, both of these cultures were derived from cultures started elsewhere in Canada). Both species were reared under standard rearing conditions. At least 21 complete wing beats were measured per species based on 2-6 wing beats per individual.

All videos of flying insects were taken at room temperature (22 ± 1°C) within a 30 × 30 × 60cm plexiglass box enclosed on three sides which functioned as a stage for filming. Prior to filming, the dorsal surface of the insect’s thorax was tethered to a straightened steel wire paperclip using heated hair removal wax. Care was taken to ensure wax did not interfere with the ability to flap. The insect was then suspended from above by a clip mounted to the top of the filming stage holding the tether. Once in the filming stage, insects were encouraged to initiate flight. This was accomplished by placing a crumpled piece of paper towel under the insect’s feet then rapidly removing the paper towel to simulate take-off or (if that failed to produce a reaction) by lightly blowing on them simulating a headwind as necessary to induce flight. The camera was placed directly behind the insect, to reflect the most probable viewing angle of a pursuing predator. High-speed video was recorded using a colour Chronos 1.4 high-speed camera (Kron Technologies, Vancouver BC, Canada) at 1770 frames per second using a Nikon 50mm (1:1.8 D) lens. Lighting was provided by a pair of 150 W halogen lamps (at ∼1m distance). Camera aperture and shutter speed were adjusted to obtain an image without motion blur between frames and with sufficient illumination to clearly distinguish the features of the subject in flight. Each captured video frame was 1024 × 768 pixels in size and the insect was framed to occupy more than 50% of the video field. Captured video sequences were stored digitally and transferred to a PC hosting software for frame-by-frame video playback (VLC media player, https://videolan.org) and single frame image analysis (ImageJ, https://imagej.net). From these videos, the wing angle at the start of the downstroke (relative to the dorsal-ventral axis of the insect), the wing beat period and projected wing area at two moments in the downstroke were measured (see below).

The wing angle at the start of the downstroke was measured from single frames using a transparent protractor placed over the computer monitor. The angle, measured from the leading edge of the wing, was relative to the dorsal-ventral axis of the animal (see Fig S1). Wing beat period (i.e. the time taken for one complete wing stroke) was measured from the frame rate used during filming and observing the number of frames elapsed between the start of each downstroke. The projected (i.e. visible) wing surface area was measured using the area measurement tool of *Image J* (open-source program, http://imagej.net, National Institutes of Health) using the known diameter of the tether wire for calibration. The tether wire diameter was chosen as it was a consistent object in the stroke plane of the insect (and in the focal plane of the camera). First, the videos were broken down into individual frames. Projected wing areas were measured from two frames during the downstroke phase of the wingbeat. The first (area1) was when the wing was approximately horizontal to the thorax (at circa half of the downstroke), and the second (area2) was just before the beginning of the upstroke (i.e. half way between mid-downstroke and the start of the upstroke). We chose to measure wing area at these phases of the downstroke to control for variation in wing beat period between beats and between species. From these frames the exposed surface area of the wing was measured (see Fig S2).

A total of 1-5 recordings (median 5) were made for each tethered individual. Seven individuals of each of seven species were observed, with the exception of the grasshopper *Melanopus femurrubrum* (six individuals observed) and the butterfly *Pieris rapae* (five individuals observed).

### Flight behaviour of species in the field

Videos of insects in the field were made using a *GoPro Hero Black 5^®^* (GoPro, San Mateo, CA). All videos were recorded at 240 frames per second. Videos were only taken under still (little wind), warm conditions (air temperatures between 18 and 25°C) when there was no precipitation. In total, 12 sympatric species were filmed, namely the grasshoppers Carolina grasshopper *Dissosteira carolina* (*n* = 12), speckled rangeland grasshopper *Arphia conspersa* (*n* = 11), *Melanoplus bivittatus* (*n* = 8) *M. femurrubrum* (*n* = 9), *M. sanguinipes* (*n* = 4) and butterflies, namely the mourning cloak *Nymphalis antiopa* (*n* = 11), monarch *Danaus plexippus* (*n* = 7), species of blue *Lycaena* spp. (*n* = 10), clouded sulphur *Colias philodice* (*n* = 13), forest tent caterpillar moth *Malacosoma disstria* (*n* = 7), least skipper *Ancyloxypha numitor* (*n* = 6) and cabbage white *Pieris rapae* (*n* =11). With the exception of *A. conspersa*, all species were filmed from September 2017 to April 2021 at sites around Ottawa, Ontario, Canada. Adult *A. conspersa* were filmed in July 2020 at sites in Corner Brook, Newfoundland, Canada. All settled insects were approached at a standard rate of approximately one pace per second by the same experimenter (KL-H). Once the targeted insect initiated its flight they were followed and recorded for as long as possible or until landing. Measurements of flight duration and proportion of time gliding were extracted from the videos recorded in the field as follows.

The duration of flight was based on the amount of flight time (in seconds) that was recorded within a video. If the entire flight was captured on video, then the flight duration was the time from take-off until landing. If at any point in the video the insect flew out of frame, the flight duration was considered to be at least the length of time recorded in the video (right censored, see statistical methods). Gliding intervals describe the periods of time (in seconds) within flight when wing beating is suspended with the wings held outstretched. Since orthoptera and lepidoptera beat their wings many times per second, the periods of time in which the wings are outstretched were unambiguous. The initiation and end of gliding intervals were recorded, and these differences summed over the entire time the individual was observed. The ratio of the sum gliding time of an individual to its total flight duration represents the proportion of flight time it spent gliding.

## Statistical Methods

All analyses were conducted in R [27] (version 4.0.3, “Bunny-Wunnies Freak Out”).

### Wing beat characteristics of species in the laboratory

To ensure that each individual provided a single multivariate data point, we began by calculating the mean wing beat period, angle, area1 and area2 per individual observed in the flight stage. We then rescaled each of the flight characteristics of individuals by subtracting the overall sample mean and dividing by the sample standard deviation, so that a unit change in each flight variable is expressed on the same scale (mean 0 and standardized deviation 1). Given that all angles were between 0 and 90 degrees, angular transformations were unnecessary. To reduce these continuous data and visualize the extent to which flight characteristics cluster according to species, we applied Principal Component Analysis (PCA) to the standardized flight characteristics of individuals (i.e. via the correlation matrix). Before drawing inferences from our PCA, we confirmed that its underlying assumptions were reasonable in that we rejected the null hypothesis of no-collinearity among predictor variables (Bartlett’s Sphericity test) and confirmed reasonable sampling adequacy of our approach based on the computed Kaiser-Meyer-Olkin (KMO) value. Our PCA was conducted using the prcomp function (default in R). Bartlett’s Sphericity test was conducted using the cortest.bartlett function, while the KMO value was computed using the paf function, both found in the rela package [28].

To determine the extent to which the flight behaviour of individuals is characteristic of the species to which they belong, we fitted a multinomial log-linear model via a neural network using the multinom function of the nnet package [29]. This multinomial model simply had the species as the response and the four principal component measures of individuals as continuous predictors. On fitting the model, we generated a confusion matrix that compared the actual and predicted species identity of each individual observed. Likewise, to quantify the dissimilarity of the flight characteristics of the different species in the wind tunnel we calculated the Mahalanobis distance between the group mean flight characteristics of each species, based on all four possible principal components (PC1-PC4) using the pairwise.mahalanobis function of the HDMD package [30].

### Flight behaviour of species in the field

The flying behaviour of grasshoppers and butterflies had to be clearly seen as they were being recorded, and so they were filmed over a range of flight times. While observations began when an insect took to the air, only 35% of individuals (largely grasshoppers) were seen to land. Thus, many of the flight durations of individuals that we recorded were “right censored” in that we can only infer that they were airborne for at least the period observed. Accordingly, we fitted Kaplan-Meir survival (time to event) models to our flight duration data which allow us to take this right censoring into account. Model fitting was via the survfit function of the survival package [31]. To obtain a quantitative estimate at the rate at which an individual settles, we separately fitted a simple intercept-only exponential model to the flight duration data of each species using the survreg function of the survival package [31]. The hazard rate (i.e. the instantaneous rate of settling) is readily calculated as the exponent of the estimated intercept of the fitted model, first multiplied by -1 (i.e. exp(-intercept)). To compare the time to settle trajectories of the different insect species we used a (non-parametric) log-rank test. To compare the mean proportion of time individuals of a given species spent gliding over the period they were monitored we used non-parametric Kruskal-Wallis test. Since our aim was to directly investigate species differences in flight behaviour rather than trait associations, we did not take the lack of independence due to shared phylogeny into account.

## Results

### Colour comparison of D. carolina and N. antiopa

The visual modelling assuming blue tit vision suggests that both the dark and light regions of *D. carolina* and *N. antiopa* wings are not distinguishable in terms of their lightness (achromatic JND < 1; **Table 1**). Chromatically, the dark regions of the two species were likely to be indistinguishable (chromatic JND ≈ 1), but the light wing regions were readily discriminable.

**Table 1.** The results of the colour discrimination model comparing the wings of *D. carolina* and *N. antiopa*. Two specimens of each species were used. JND refers to just noticeable difference.

| Viewing angle | Compared region | Chromatic JND | Achromatic JND |
| --- | --- | --- | --- |
| Dorsal | Dark region | 0.98 | 0.25 |
|  | Light region | 13.09 | 0.35 |
| Ventral | Dark region | 1.02 | 0.19 |
|  | Light region | 9.06 | 0.27 |

### Wing beat characteristics of species in the laboratory

Fig 2 shows the underlying distributions of our continuous flight variables which showed high degrees of (linear) association, notably among angle and wing beat period as well as visible wing surface area (from measurements of area1 and area2). This collinearity was confirmed via Bartlett’s test (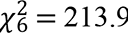, P < 0.001) while the KMO value was 0.59, indicating reasonable sampling adequacy for PCA.

**Fig 2.**
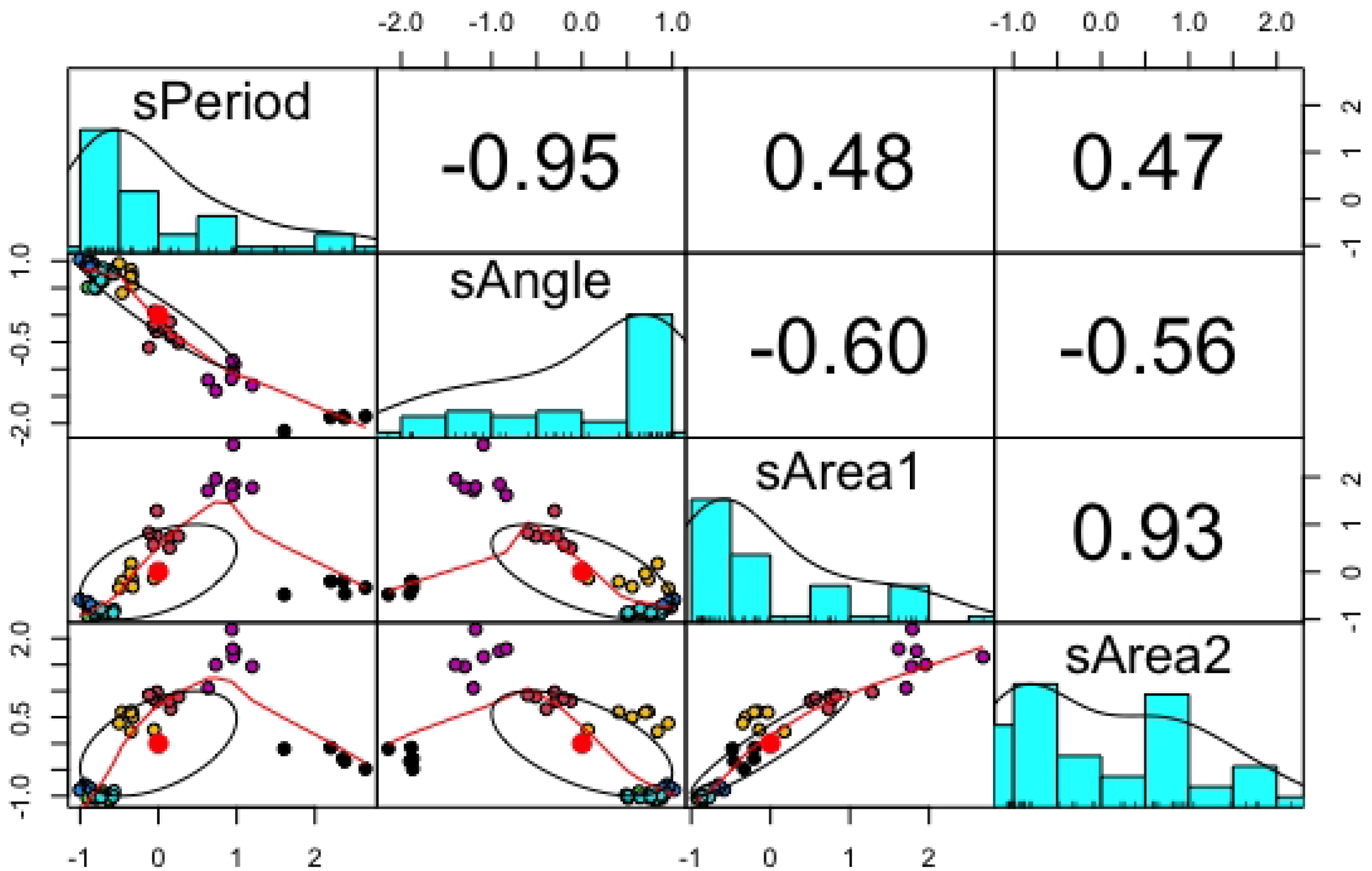
The distributions, bivariate scatter plots and Pearson correlations among the (re-scaled) mean flight characteristics of individuals from the different species. The “s” prefix refers to the fact that the variables had been standardized. The wing beat period and angle of individuals were highly negatively correlated, while area1 and area2 were highly positively correlated. The red dots refer to the sample means of the standardized distribution (0 given the re-scaling) while different colours are used to depict species: *Dissosteira carolina*, red; *Locusta migratoria*, yellow; *Melanoplus sanguinipes*, green; *Melanopus bivittatus*, blue; *Melanopus femurrubrum*, turquoise; *Nymphalis antiopa*, purple; *Pieris rapae,* black.

On conducting the PCA, the first principal component (PC1) explained 75.0% of the variance while the second (PC2) explained an additional 22.3% of the variance. With a cumulative proportion of 97.3% of the variance explained by the first two principal components (and only 1.87% explained by PC3) then most of the inter-individual variation in flight characteristics can be visualized by plotting these two orthogonal components. Fig 3 shows the principal component plot for the 46 tethered individuals in our experiment. The multivariate flight characteristics of individuals were such that they fall into clearly defined species clusters, with the exception of the three *Melanopus* grasshopper species which show considerable overlap in their 95% elliptical iso-contours.

**Fig 3.**
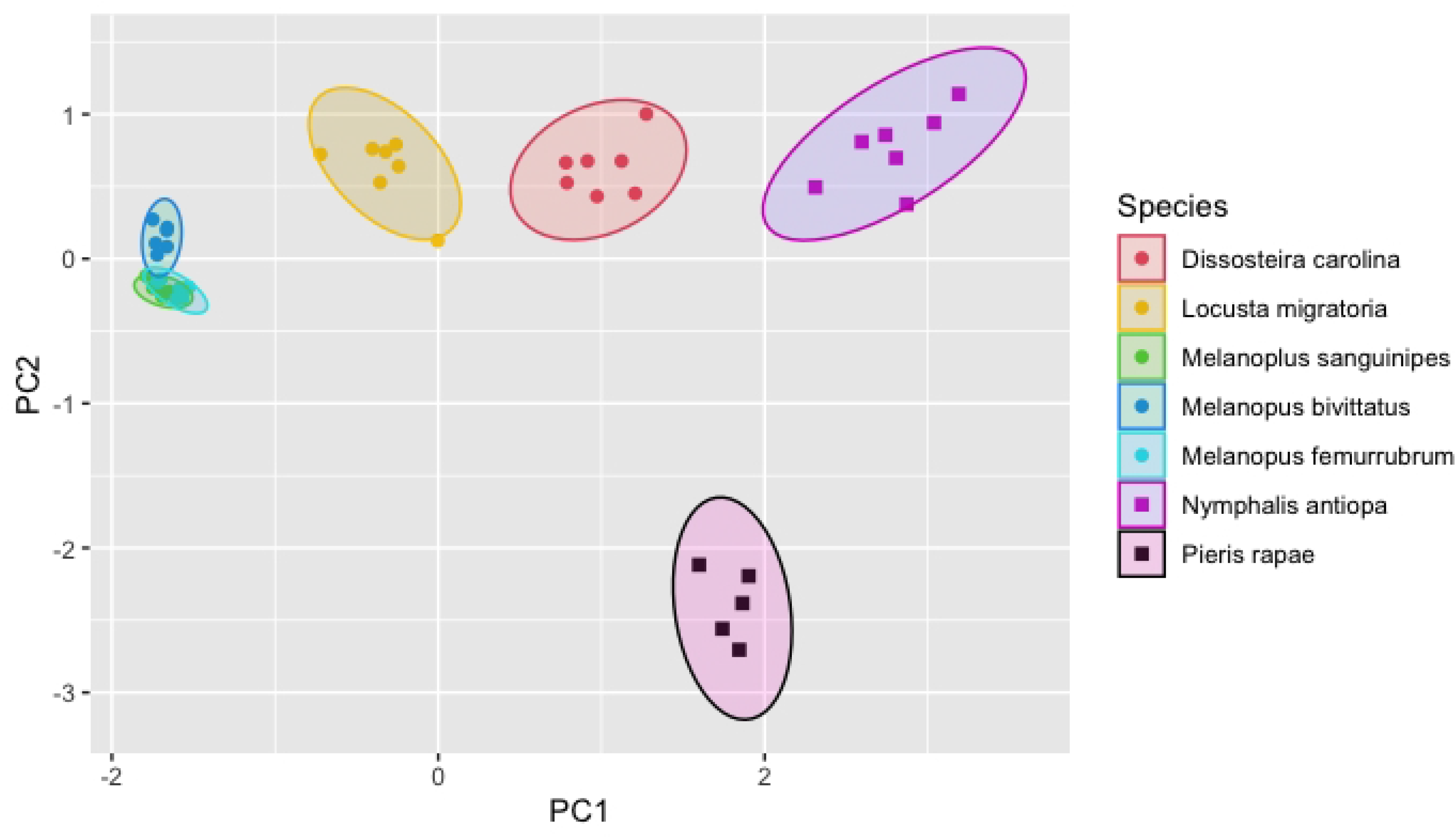
Plot of the flight characteristics of the 46 tethered individuals observed in the laboratory, based on orthogonal (PC1 and PC2) linear combinations of their wing beat period, angle, area1 and area2. The species to which each individual belongs is depicted by their colour while their insect order is depicted by shape (circle, Orthoptera; square, Lepidoptera). The filled ellipses show the 95% confidence interval isoclines for the principal components for each species, assuming multivariate normality within each species cluster.

The confusion matrix (**Table 2**) generated by the fit of our neural net model to the principal components confirms the above interpretation, suggesting that most species could be readily distinguished on the basis of their multivariate flight characteristics, the exception being *M. sanguinipes* and *M. femurrubrum*. From visual inspection of the principal component plot, the (*D. carolina*, red circle) has the closest flight behaviour to the mourning cloak butterfly (*N. antiopa*) of any species investigated, including the other butterfly species in our sample (*P. rapae*). The estimated dissimilarity matrix (**Table 3**) of species further supports this view, with the smallest Mahalanobis distance (0.07) reported between *M. sanguinipes* and *M. femurrubrum*. Moreover, while the Carolina grasshopper (*D. carolina*) and mourning cloak butterfly (*N. antiopa*) could be quantitatively discriminated on the basis of their flight characteristics (**Table 3**), the overall closest match to *N. antiopa* was *D.carolina* (Mahalanobis distance of 1.81, **Table 3**).

**Table 2.** The species classification of individuals based on fitting a neural net model that distinguished between species using the principal components of their flight characteristics observed in the laboratory. The species to which individuals are predicted to belong are compared with the species to which they actually belong (*Dissosteira carolina*, *D.c*; *Locusta migratoria*, *L.m*; *Melanoplus sanguinipes*, *M.s*; *Melanopus bivittatus*, *M.b*; *Melanopus femurrubrum*, *M.f*; *Nymphalis antiopa*, *N. a*; *Pieris rapae, P.r*). Only *M.f* and *M.s* were confused based on their multivariate flight characteristics, and even here the confusion was partial.

|  |  | Actual species |  |  |  |  |  |  |
| --- | --- | --- | --- | --- | --- | --- | --- | --- |
|  |  | *D.c* | *L.m* | *M.b* | *M.f* | *M.s* | *N.a* | *P.r* |
| Predicted species | *D.c* | 7 | 0 | 0 | 0 | 0 | 0 | 0 |
|  | *L.m* | 0 | 7 | 0 | 0 | 0 | 0 | 0 |
|  | *M.b* | 0 | 0 | 6 | 0 | 0 | 0 | 0 |
|  | *M.f* | 0 | 0 | 0 | 5 | 1 | 0 | 0 |
|  | *M.s* | 0 | 0 | 0 | 2 | 6 | 0 | 0 |
|  | *N.a* | 0 | 0 | 0 | 0 | 0 | 7 | 0 |
|  | *P.r* | 0 | 0 | 0 | 0 | 0 | 0 | 5 |

**Table 3.**
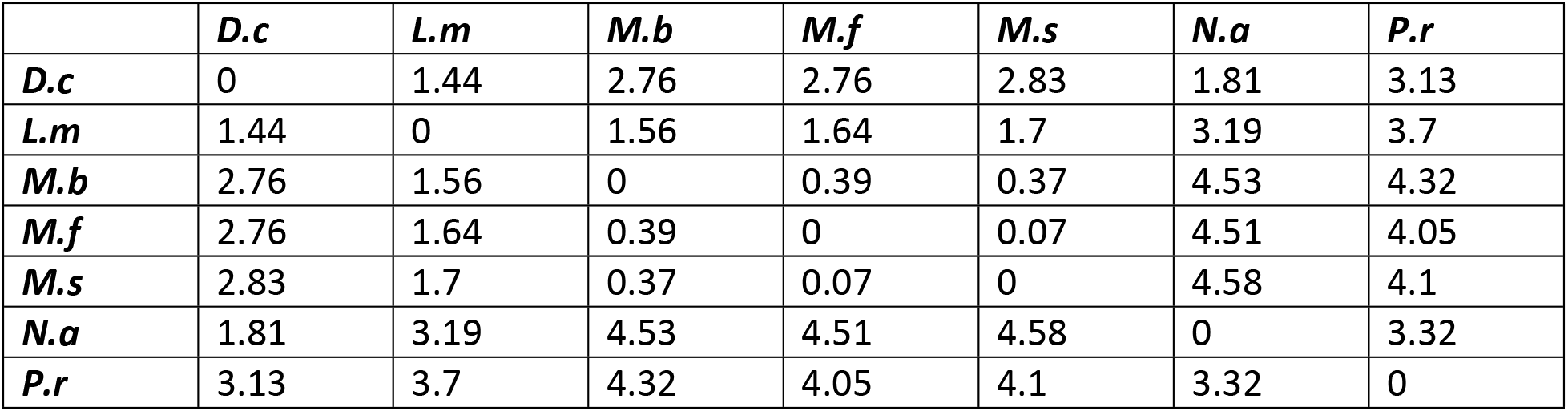
The Mahalanobis distances between the mean principal components of each species derived from their flight behaviour in the lab. Abbreviations: *Dissosteira carolina*, *D.c*; *Locusta migratoria*, *L.m*; *Melanoplus sanguinipes*, *M.s*; *Melanopus bivittatus*, *M.b*; *Melanopus femurrubrum*, *M.f*; *Nymphalis antiopa*, *N. a*; *Pieris rapae, P.r*. These distances can be considered a measure of their overall behavioral similarity, with shorter distances reflecting a closer similarity. The closest species to *N. a* (i.e. lowest dissimilarity) was *D.c*, while *D.c* was closest in flight behaviour to *L.m* and *N.a*.

|  | *D.c* | *L.m* | *M.b* | *M.f* | *M.s* | *N.a* | *P.r* |
| --- | --- | --- | --- | --- | --- | --- | --- |
| *D.c* | 0 | 1.44 | 2.76 | 2.76 | 2.83 | 1.81 | 3.13 |
| *L.m* | 1.44 | 0 | 1.56 | 1.64 | 1.7 | 3.19 | 3.7 |
| *M.b* | 2.76 | 1.56 | 0 | 0.39 | 0.37 | 4.53 | 4.32 |
| *M.f* | 2.76 | 1.64 | 0.39 | 0 | 0.07 | 4.51 | 4.05 |
| *M.s* | 2.83 | 1.7 | 0.37 | 0.07 | 0 | 4.58 | 4.1 |
| *N.a* | 1.81 | 3.19 | 4.53 | 4.51 | 4.58 | 0 | 3.32 |
| *P.r* | 3.13 | 3.7 | 4.32 | 4.05 | 4.1 | 3.32 | 0 |

### Flight behaviour of species in the field

Fig 4a,b shows the fitted “survival” (i.e. the duration from the flight initiation until the settling) curves of the Orthoptera (5 species) and Lepidoptera (7 species) in our recordings. While the vast majority of Orthoptera settled within 120 seconds; of the Lepidoptera only the mourning cloak butterfly (*N. a*) and least skipper (*An. n*) tended to come to rest within this time period. The estimated hazard rates of these two Lepidoptera species were correspondingly high (**Table 4**). Of all the Orthoptera species observed, the speckle-winged rangeland grasshopper (*A. c*) and the Carolina grasshopper (*D. c*) exhibited the longest time in flight (Fig 4a and Table 4). While the settling curves differed among Orthoptera species (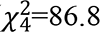, P < 0.001) and among Lepidoptera species (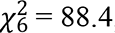, P < 0.001) there was no significant difference in the settling rate of the Carolina grasshopper and mourning cloak butterfly (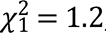, *P* = 0.30, Fig S3).

**Fig 4a,b.**
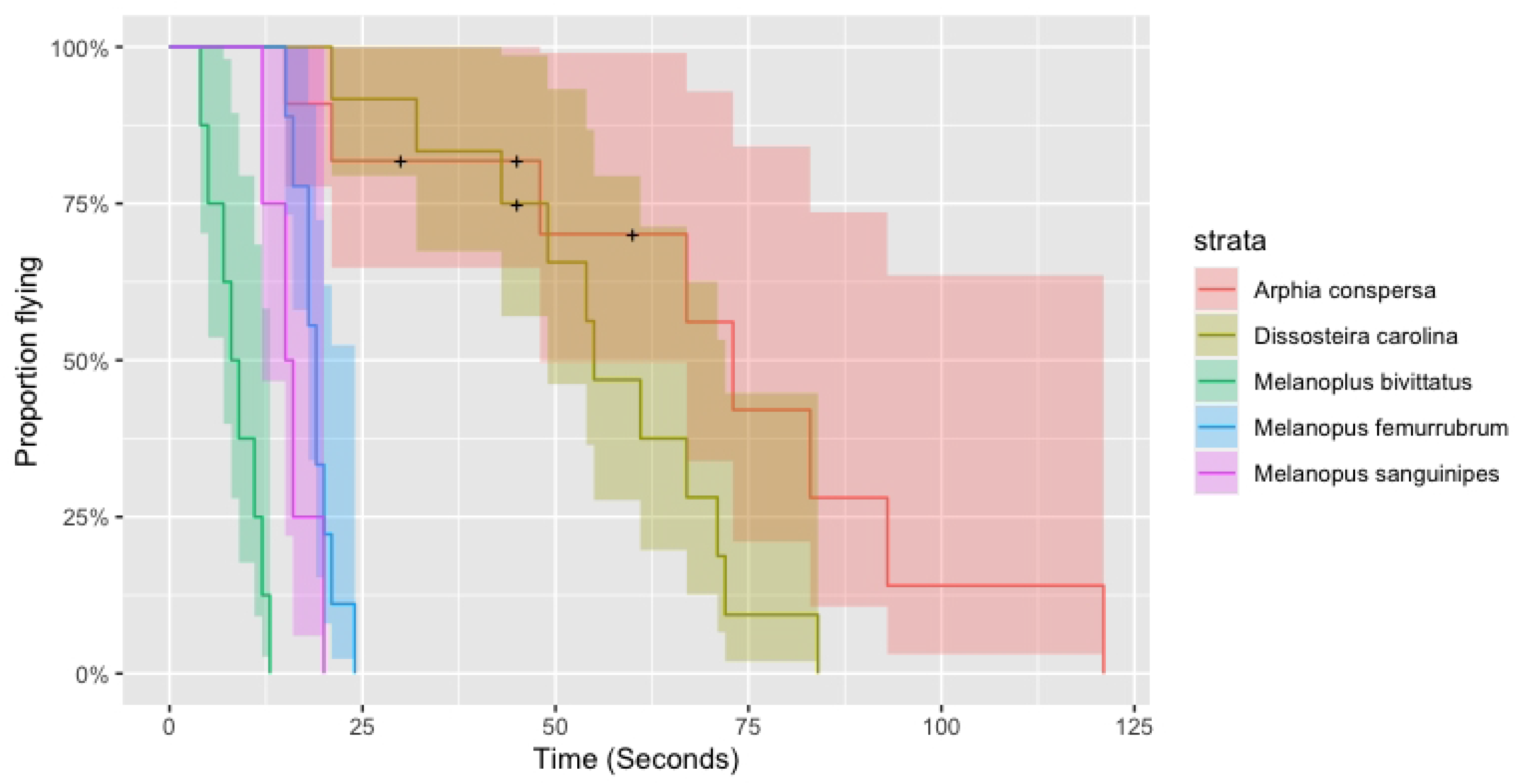
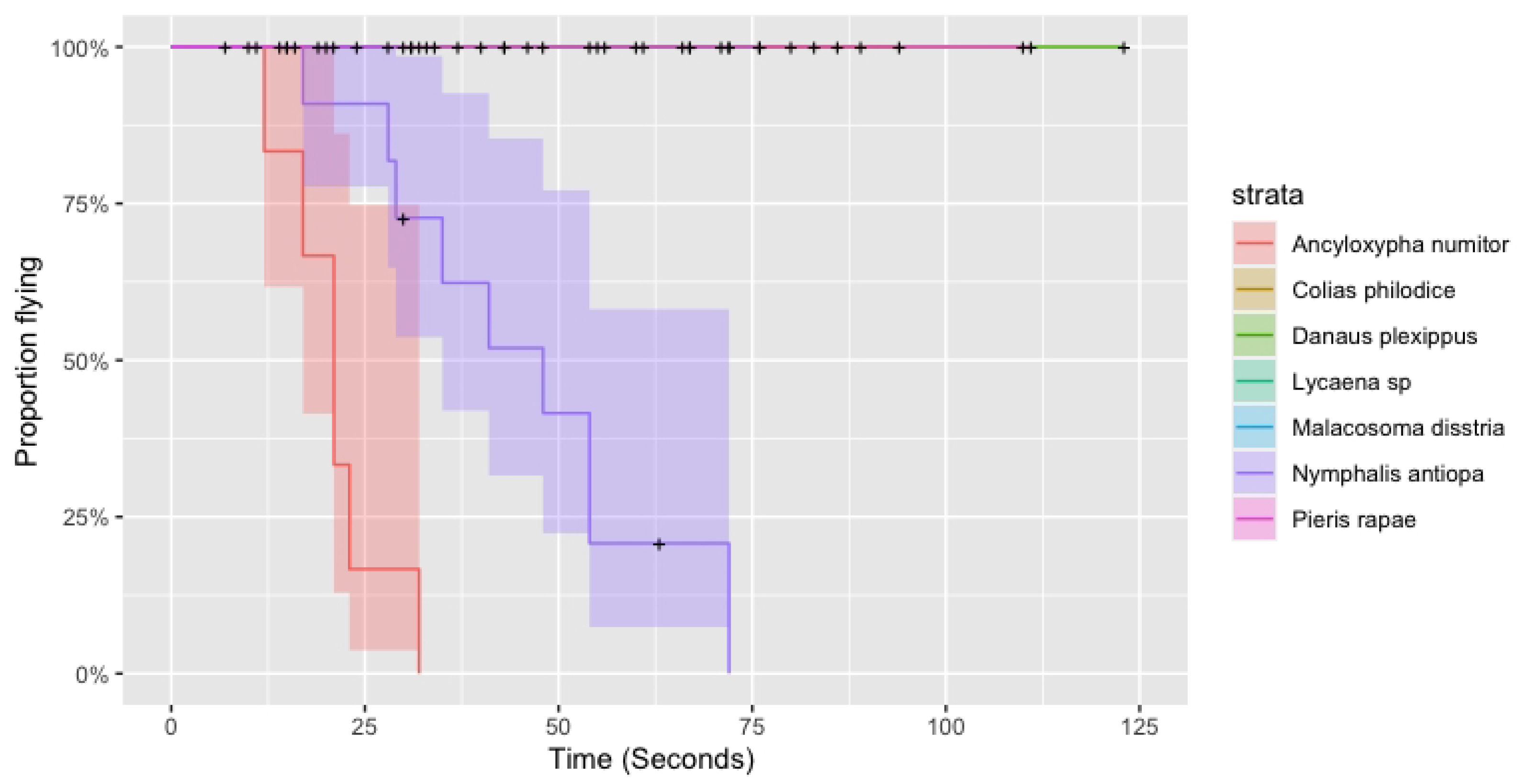
Fit of survival curves to the flight durations of (a) Orthoptera and (b) Lepidoptera. Legend abbreviations: the crosses refer to right-censused individuals (settling not observed) while shading shows the 90% confidence intervals. The majority of Orthoptera observed settled within two minutes: of all the Orthoptera species observed, the speckle-winged rangeland grasshopper (*Arphia conspersa*) and the Carolina grasshopper (*Dissoteira carolina*) exhibited the longest time in flight. By contrast most individuals of the lepidopteran species did not settle within two minutes, so the proportion flying remained at 100%. Of the Lepidoptera only the mourning cloak butterfly (*Nymphalis antiopa*) and least skipper (*Ancyloxpha numitor*) settled within two minutes.

**Table 4.** Estimates of the rate of settling of the different insect species based on fitting an exponential model of decay to their observed time before settling (including individuals that were censused). Of the butterflies, only the least skipper (*Ancyloxypha numitor*) and mourning cloak (*Nymphalis antiopa*) had settling rates that were comparable with orthopterans we observed. The morning cloak *Nymphalis antiopa* (Lepidoptera) and Carolina grasshopper *Dissosteira carolina* (Orthoptera) settled at very similar rates.

| Species | Intercept | SE | Hazard | Taxon |
| --- | --- | --- | --- | --- |
| *Melanoplus bivittatus* | 2.15 | 0.354 | 0.116 | Orthoptera |
| *Melanoplus sanguinipes* | 2.76 | 0.500 | 0.063 | Orthoptera |
| *Melanopus femurrubrum* | 2.94 | 0.333 | 0.053 | Orthoptera |
| *Ancyloxypha numitor* | 3.04 | 0.408 | 0.048 | Lepidoptera |
| *Nymphalis antiopa* | 3.96 | 0.333 | 0.019 | Lepidoptera |
| *Dissosteira carolina* | 4.09 | 0.302 | 0.017 | Orthoptera |
| *Arphia conspersa* | 4.41 | 0.354 | 0.012 | Orthoptera |
| *Danaus plexippus* | 6.74 | 1.000 | 0.001 | Lepidoptera |
| *Malacosoma disstria* | 34.65 | 3178309 | 0.000 | Lepidoptera |
| *Lycaena sp* | 35.63 | 2624107 | 0.000 | Lepidoptera |
| *Colias philodice* | 35.71 | 2306698 | 0.000 | Lepidoptera |
| *Pieris rapae* | 36.1 | 2590841 | 0.000 | Lepidoptera |

Fig 5a,b shows the distribution of the total time insects of a given species spent gliding as a proportion of the time they were observed. Of the Orthoptera (Fig 5a), only the Carolina grasshopper consistently spent time gliding (7 of 12 individuals). The arithmetic mean proportion of time this species was observed gliding was 0.11 (sd 0.11) while the overall (weighted) mean proportion of time spent gliding was 82/564 = 0.13. The only other orthopteran observed gliding was the speckle-winged rangeland grasshopper (*Arphia conspersa*) with 1 of 11 individuals recorded gliding, with an arithmetic mean proportion of time spent gliding of 0.01 (sd 0.03) and a weighted mean of 3/656 i.e. 0.005. Given the variability among species, we can reject the null hypothesis that there was no systematic variation in gliding rates among orthopteran species (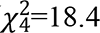, P=0.001). Of the Lepidoptera (Fig 5b), only the mourning cloak (8 of 11 individuals) and monarch (7 of 7 individuals) butterflies exhibited gliding behaviour over the time recorded. The arithmetic mean proportion of time the mourning cloak was observed gliding was 0.26 (sd 0.20) while weighted mean proportion of time spent gliding was 148/471 i.e. 0.31. The arithmetic mean proportion of time the monarch was observed gliding was 0.12 (sd 0.08) while weighted mean proportion of time was 87/842 = 0.10. Given the variability, we can reject the null hypothesis that there was no systematic variation in gliding rates among butterfly species (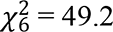, P < 0.001).

**Fig 5a,b.**
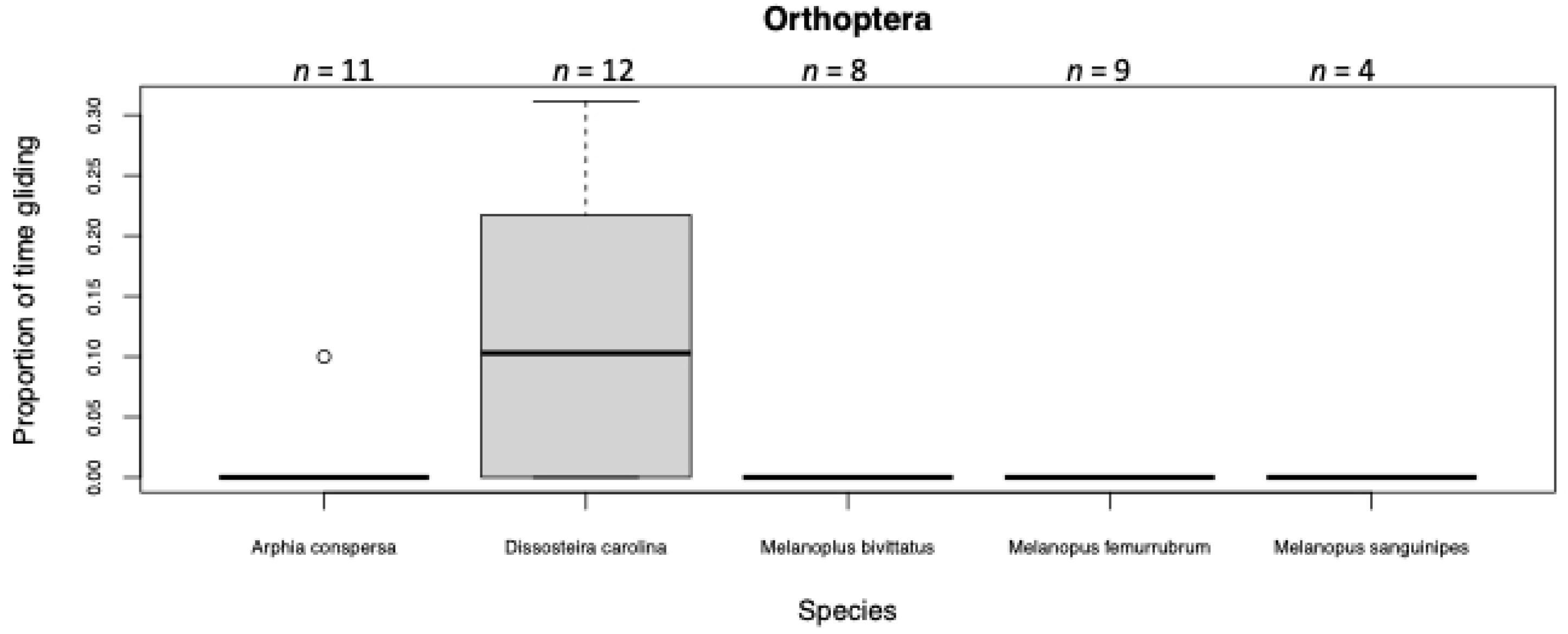
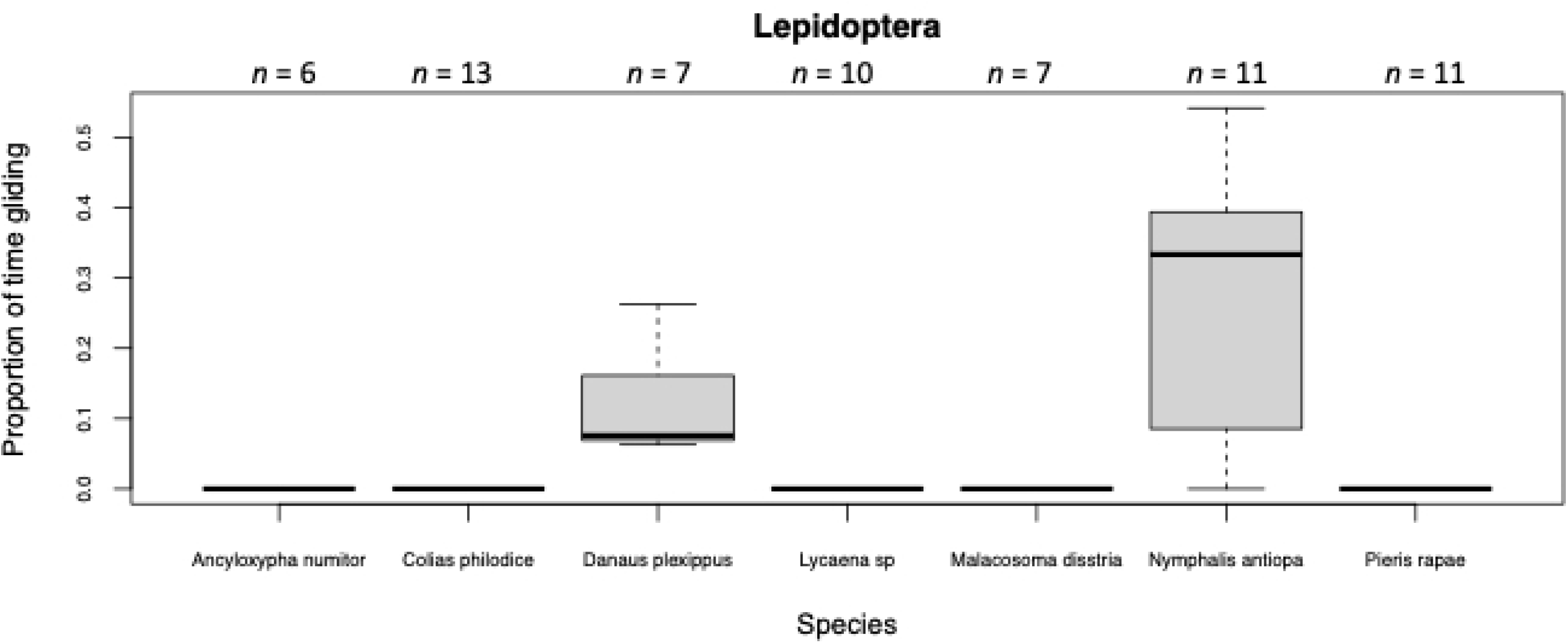
Box and whisker plots proportion of time individuals of a given orthopteran species (Fig 5a) or lepidopteran species (Fig 5b) were observed gliding as they flew. The dark line shows the median, the boxes show the interquartile range, while the whiskers show the extremes up to an additional 1.5 of the interquartile range. Points are outliers beyond this range.

In sum, the only orthopteran species that spent a non-negligible amount of time gliding was the Carolina grasshopper, while the butterfly that glided the most in our species sample was the mourning cloak. However, the distributions of the glide rates of individuals differed significantly between these two species (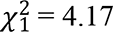, P < 0.041) – see Fig S3, with the mourning cloak spending a slightly higher proportion of its time gliding (Fig S4).

## Discussion

To human eyes, the hind wings of *D. carolina* appear similar in both colour and pattern to those of *N. antiopa.* Indeed, given their spectral properties, the results of our visual modeling suggest that avian predators would be unable to detect a difference in the hue of the dark brown surfaces on the wings of the two species. These dark surfaces cover a high proportion of the hind wing surface areas of *D. carolina* and *N. antiopa*, and likely contributes to their substantial overall perceived visual resemblance. The light region on the anterior wing margins of the two species would in principle be chromatically (but not achromatically) distinguishable to a bird, but this may not be a salient distinguishing feature. Indeed, it has been demonstrated that the flash-lag effect causes vertebrates to perceive the margins of objects moving at high velocity to chromatically blend with both the center of the moving object and its background [32]. Therefore, the coloration on the wing margins of *D. carolina* may be of less consequence to the perceived appearance of this species in flight.

While a degree of similarity in colour pattern may facilitate mimicry, coloration is clearly not the only feature used by birds when distinguishing between different categories of stimuli [33]. Motion plays an important role in shaping the way observers detect and discriminate among targets [34, 35]. Our laboratory data suggest that, of all the species investigated, the Carolina grasshopper is the species most similar to the mourning cloak in terms of its flight characteristics among the co-occurring orthopterans. Likewise, our field data suggest that of all the species investigated, the Carolina grasshopper is most similar to the mourning cloak in terms of settling rate and most similar in terms of gliding rate. The similarity of *D. carolina* and *N. antiopa* not just in terms of colour pattern but also in terms of flight behaviour provides reasonably strong evidence for a mimetic relationship because the more independent dimensions the similarity holds between these distantly related species, the more likely it has not arisen by chance.

Despite the similarities in flight and colour, if one looks closely enough at the behaviour and/or appearance of mimics or models in any system then one can always distinguish them [36]. In our case, the neural net could distinguish between species based on unique combinations of their wing beat duration, angle, area1 and area2. Birds are known to have ultra-rapid (high-flicker fusion frequency) vision [37] capable of detecting changes in fast-moving prey, but it is unclear whether they could detect the differences in the flight characteristics we have uncovered. Even if predators could in principle distinguish between species, one might wonder whether they would ever be motivated to classify prey to species level if all they wish to ascertain if they are worth pursuing. As always therefore, experiments with real predators are necessary to fully test the hypothesis that they are in a mimetic relationship.

The common colour pattern and behaviour of the Carolina grasshopper and mourning cloak butterfly is not particularly surprising, since selection for mimetic resemblance frequently extends beyond appearance to behavioral attributes, including the way the mimic moves [38, 39]. For example, it has been recently shown that the wingbeat patterns of mimetic forms of the polymorphic swallowtail butterfly *Papilio polytes* match those of its unpalatable model *Pachliopta aristolochiae* it resembles [40]. Likewise, the salticid spider *Myrmarachne formicaria* not only looks like *Myrmarachne* ants in appearance, but also walks like them [41]. Similarly, the common drone fly (*Eristalis tenax*) flies in a manner similar to the honeybee (*Apis mellifera*) it morphologically resembles [42]. The Bornean gliding lizard, *Draco cornutus* have gliding membranes that closely match the colours of local freshly falling leaves, but their gliding behaviour is also similar to one of many ways that leaves fall in the breeze [43].

If the Carolina grasshopper and mourning cloak butterfly are indeed engaged in a mimetic relationship, then a natural question to ask is why the mimicry has arisen. Online field guides suggest that both the Carolina grasshopper and mourning cloak butterfly are palatable to consume, in that they are both attacked by avian predators (Carolina grasshopper [20], mourning cloak: [44]). In Ontario, the morning cloak butterfly emerges as an adult in the early spring months before the Carolina grasshopper emerges in mid to late summer [45]. Since experience of the model is typically necessary for predators to select for mimicry, the later emergence of the Carolina grasshopper suggests it has evolved to resemble the morning cloak, and not *vice versa*. It is perhaps not coincidental that the mourning cloak butterfly spends a relatively low proportion of its time flying compared to most butterflies, rendering it a relatively easy target for mimicry by grasshoppers with low flight durations.

In our introduction we discussed the possibility that the display causes predators to adopt a sub-optimal search strategy, giving up sooner when it does not find the signaller that appears to have settled [1, 3] (see also our SI). However, it is not the only explanation for the evolved mimicry. Like many butterflies, the mourning cloak is relatively hard to catch, at least by humans. One possibility therefore is that the slower *D. carolina* has evolved to resemble a harder to catch *N. antiopa*, which would make it an example of (Batesian) “evasive mimicry” [46-48]. Indeed, the speckled rangeland grasshopper *Arphia conspersa* (which was notable for its long time in flight and ability to glide in our field study) similarly appears to resemble the alfalfa butterfly *Colias eurytheme*/*C. philodice* not only in hind wing colour pattern but also in flight behaviour [49]. In this case the grasshopper was demonstrated to be easier to catch (by a human) than the butterfly, leading Balgooyen [49] to conclude that the grasshopper was an evasive Batesian mimic. Loeffler-Henry & Sherratt [50] recently argued for yet another form of mimicry, which they dubbed “mutualistic deceptive mimicry” between profitable species that are caught in different ways by predators. In these instances, there may be selection on both prey species to resemble one another because it increases the chance that a predator will use an inappropriate (and hence inefficient) attack strategy to catch them.

It is important to note that mimicry can benefit a signaller at multiple stages through the predation sequence, so the above explanations are not mutually exclusive. For example, the speckled rangeland grasshopper described above is highly cryptic when at rest (“virtually undetectable” [49]). So, even if the resemblance arises in part as a consequence of Batesian evasive mimicry (pursuit deterrence), then this does not preclude the possibility that the mimetic flash behaviour maintains the chances of remaining undetected after the prey has settled (a search deterrence). Our Bayesian search theory model (SI Appendix A) readily accounts for both forms of mimicry in the same underlying framework. Thus, if models are common and relatively unprofitable to pursue then predators will not be motivated to search for the model/mimic when it settles. However, even if the predator does pursue the fleeing prey, it may give up searching sooner as a consequence of the mimicry.

## Supporting Information

**Appendix A: A Bayesian model of flash display mimicry.**

**Fig S1. Examples of maximal stroke plane angle measurement.** a. *Locusta migratoria*, b. *Nymphalis antiopa*.

**Fig S2. Examples of projected wing area measurement.** a. *Nymphalis antiopa* Area1, b. *N. antiopa* Area2, c. *Locusta migratoria* Area1, d. *L. migratoria* Area2.

**Fig S3. Fit of survival curves to the flight durations of the Carolina grasshopper *Dissosteira Carolina* and the mourning cloak, *Nymphalis antiopa*.** The crosses refer to right-censused individuals (settling not seen) while the shading shows the 90% confidence intervals. Overall, the two species had similar rates of settling that were statistically indistinguishable based on the sample data.

**Fig S4. Box and whisker plot of the proportion of flying time individuals of the Carolina grasshopper *Dissosteira carolina* and the mourning cloak, *Nymphalis antiopa* spent gliding.** Although the Carolina grasshopper exhibited an unusually high propensity to glide for an orthopteran, it was still significantly less than the overall rate of gliding of the mourning cloak.

## Acknowledgments

We are grateful to Kate Derbyshire and Kiki Malott who collected mourning cloak caterpillars that were raised to adulthood and used in the flight stage. We also thank Jane Allision who kindly allowed us to use *Melanoplus sanguinipes* from her laboratory culture in the flight stage experiments. KL-H is supported by an NSERC Postdoctoral Fellowship. CK is supported by National Research Foundation of Korea (NRF-2019R1C1C1002466). TNS is supported by an NSERC Discovery Grant.

## Author contributions

**Conceived and designed the experiments:** TNS, KLH, CK, JD.

**Performed the experiments:** KLH, CK.

**Analyzed the data:** TNS, CK.

**Mathematical modelling:** TNS.

**Wrote the paper:** TNS, KLH, CK, JD.

